# Exploring Pharmacogenetic Factors Influencing Hydroxyurea Response in Tanzanian Sickle Cell Disease Patients: A Genomic Medicine Approach

**DOI:** 10.1101/2024.05.23.595488

**Authors:** Siana Nkya, Collin Nzunda, Emmanuel Saukiwa, Frida Kaywanga, Eliud Buchard, David Solomon, Heavenlight Christopher, Doreen Ngowi, Julieth Johansen, Florence Urio, Josephine Mgaya, Christina Kindole, Mbonea Yonazi, Salman Karim, Mohamed Zahir Alimohamed, Raphael Z. Sangeda, Clara Chamba, Collet Dandara, Enrico Novelli, Emile R. Chimusa, Julie Makani

## Abstract

Sickle cell disease (SCD) continues to pose a significant public health challenge, particularly in sub-Saharan Africa. Despite its discovery over a century ago, the progress in developing and accessing effective interventions has been notably restricted. Currently, hydroxyurea stands as the primary drug in widespread use, and has been associated with elevated levels of fetal hemoglobin (HbF) and enhanced clinical outcomes. Notably, a substantial proportion, up to 30%, of patients do not exhibit a positive response to hydroxyurea treatment. There is compelling evidence suggesting that genetic factors play a crucial role in influencing the effectiveness of hydroxyurea. In this study, we present findings on the investigation of genetic variants influencing hydroxyurea response in 13 genetic loci associated with HbF synthesis and hydroxyurea drug metabolism focusing on *MYB*, *HBB*, *HBG1*, *HBG2*, *BCL11A*, *KLF10*, *HAO2*, *NOS1*, *ARG2*, *SAR1A*, *CYP2C9*, *CYP2E1*. We report remarkable genetic associations with *CYP2C9*, *CYP2E1, KLF10*, *BCL11A*, *ARG2*, *HBG1*, *SAR1A*, *MYB*, and *NOS1* loci with hydroxyurea response. We also highlight associated pathway’s enrichment and gene-gene interactions analysis in the context of hydroxyurea treatment response.

## Introduction

Hydroxyurea (HU) is the only FDA-approved drug for sickle cell disease (SCD) and a major intervention for the disease. It was first approved for use in SCD in 1998 [1] and has been shown to increase levels of fetal hemoglobin (HbF), which is a major disease modifier for SCD. Although the mode of action of hydroxyurea has not been completely elucidated, hydroxyurea induces gamma-globin expression that leads to increase HbF. To date, many SCD patients have received hydroxyurea treatment, and those who have responded well have experienced reduction of VOC by 50% as well as decreased need for transfusions [2,3]. However, response to HU treatment varies from one patient to another with very good, moderate and poor responders. Weight-based dosing with stepwise escalation to maximum tolerated dose (MTD) is recommended and is associated with predictable laboratory (increased HbF and total Hb levels) and clinical benefits [4]. However, this process takes 6 to 12 months to achieve and confirm treatment response based on hematological profile changes of the patients compared to their baseline levels [5]. In addition, in many African countries, the routine clinical practice is provision of a fixed HU dose due to limited accessibility to the drugs as well as monitoring of drug tolerance [6].

Hydroxyurea treatment response can be monitored/predicted at different levels including genetic, molecular, and haematological (**Supplemental_Fig_1**). While molecular and haematological profiles can only be measured post treatment, genetic factors/markers can be determined prior to treatment commencement and hence posing as the best markers for predicting drug response. Genetic variants associated with HbF induction and hydroxyurea metabolism are among factors that influence HU treatment response [7–9]. Most of the studies to investigate the genetic variants associated with hydroxyurea treatment (hydroxyurea pharmacogenomics studies) have been conducted in non-African countries [10,11]. In addition, the identified genetic variants have not been developed into pharmacogenetic tests for HU treatment. When available, hydroxyurea pharmacogenetic test will be a one-time solution for an early HU response prediction which may enhance decision of either dose escalation or the need for alternative drugs/therapy. We have investigated the genetic determinants of HU treatment response and have identified associated genetic variants in *CYP2C9*, *KLF10*, *BCL11A*, *ARG2*, *HBG1*, *SAR1A*, *MYB*, and *NOS1* that are potential predictors of hydroxyurea response.

## RESULTS

### Hematological profiles pre and post hydroxyurea treatment

This was a longitudinal study including a total of 148 individuals with sickle cell disease (SCD) in Tanzania, encompassing both genders (58% female) with the age ranging from 5 to 30 years (mean ± SD: 24 ± 11.49). The individuals were enrolled and followed up for six-month period with scheduled monthly visits.

Over a period of 6 months, 148 patients with sickle cell disease were treated with Hydroxyurea (HU) and closely monitored. During this period, notable variations were observed in the haematological indices. Specifically, slight increases were noted in Red Blood Cell count (RBC) from a median of 2.7 ± 1.05 to 2.79 ± 1.12, Hemoglobin (HB) from 7.5 ± 1.43 to 7.63 ± 2.53 and Mean Corpuscular Volume (MCV) from 82.4 ± 12.2 to 83 ± 11.2. Conversely, a decrease was observed in reticulocytes from a median of 10.9 ± 5.61 to 9.63 ± 6.87 by the third visit. We could not establish significant changes in platelets (381 ± 205 to 382 ± 172), white blood cells (12.9 ± 6.09 to 12.8 ± 6.81), and fetal hemoglobin (5.3 ± 4.61 to 5.2 ± 5.08).

However, when the patients were categorised either as poor or good responders based on HbF levels, significant differences in haematological indices were observed between the two groups. Of the 148 participants subjected to hydroxyurea (HU) treatment, sixty (60) individuals were effectively stratified into either responder or non-responder categories while the rest were treated as the ‘unclassified’ group, comprising individuals for whom clear distinctions based on HbF levels were challenging or not applicable. Two-fold increase in HBF from the baseline (first visit) were considered as responder and vice -versa. Within this classification, forty-nine (49) individuals demonstrated a favourable response, while eleven (11) were identified as non-responders or poor responders.

Our findings indicated that the effect of hydroxyurea treatment (represented by visits) on the haematological indices varied significantly (p < 0.01) between the poor and good responders. The estimated marginal means were also calculated for the interaction between respondence and visits, along with their 95% confidence intervals. Hydroxyurea treatment responders showed the highest estimated mean increases throughout the visits in HBF and MCV compared to the poor responders. On the other hand, poor responders had higher levels of WBC and RET compared to the good responders (**Figure 1**) .

**Figure 1.**
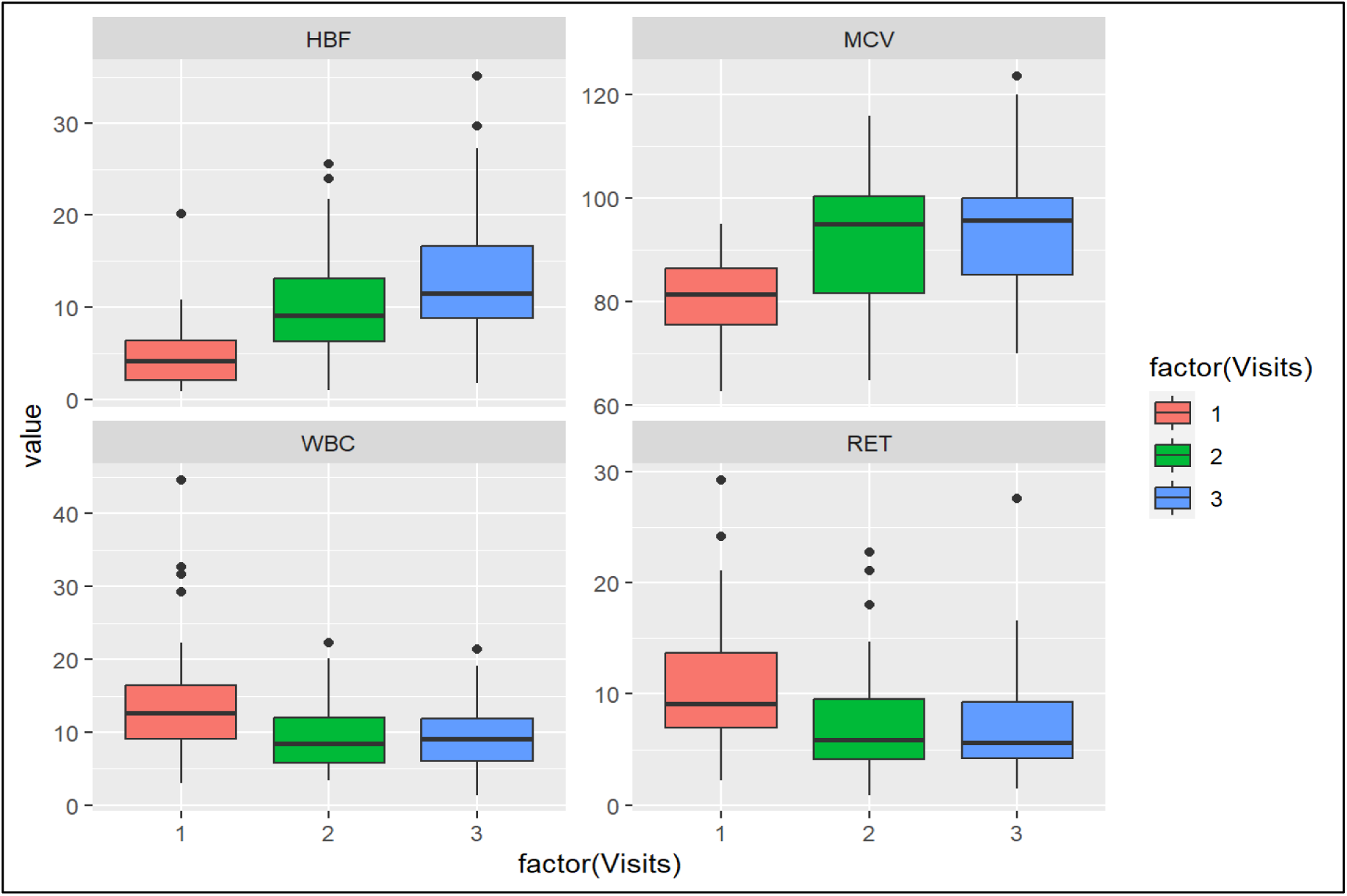
Haematological profiles of poor and good responders to hydroxyurea treatment. Good responders showed the highest estimated means increase throughout the visits in HBF and MCV while poor responders showed the highest in WBC and RET. Results have indicated that the effect of time after treatment (visits) on the dependent variables varies significantly with p-value of < 0.01 based on different levels of responding status.

### Genetic profiles for the hydroxyurea good and poor responders

The genetic analysis included a total of 49 individuals (good and poor responders). Our analysis utilised several bioinformatics and population genomics tools to leverage free hypothesis approaches such as mutation prioritizing instead of performing statistical testing model such as case/control association that require to reach a level of significance based on sample size and power, when comparing 2 groups of samples (**Supplemental_Fig_3**).

### Targeted variant discovery in hydroxyurea good responders and poor responders

A total number of 152,327 variants (10.8 % insertion, 6.7% deletion, 8.5% structural variants, 0.001% multi-nucleotide variants, and 74% SNPs) were successfully called from the targeted sequence panel (**Supplemental_Table_2A**), of which 1% and 52% were exonic, and intergenic respectively, distributed as 0.003% stop loss, 0.026% stop gain, 0.5% synonymous, and 0.36% non-synonymous, 0.02% splice site variants in dataset. **Supplemental_Fig_2B** illustrates the quality control of the sequence alignment data.

### Variant prioritization, mutation burden and proportion of pathogenic variants

Among hydroxyurea good responders and poor responders, we detected significant differences in the burden of non-synonymous, function-altering variants (**Table 1**). Potential pathogenic Single Nucleotide Polymorphism (SNPs) were identified by selecting variant predictions of deleteriousness (**Table 1**) from at least 10 out of 14 predictive tools using ANNOVAR (**Materials and Methods**). Therefore, a trimmed list of nine Single Nucleotide Variation (SNVs) that were classified as “damaging” by FATHHM is presented (**Table 1**). The most deleterious mutations were found within the *HAO2* [MIM: 605176], *BCL11A* [MIM: 606557], *WAC* [MIM: 615049], *CYP2C9* [MIM: 601130], *CYP2E1* [MIN: 124040], *MYB* [MIM: 189990], *HBG1* [MIM:142200], *NOS1* [MIM 163731] and *ARG2* [MIM 107830] variants (**Table 1**, **Figure 2A**) of which four variants were found only in the “responders” group (*MYB* [MIM: 189990], *HBG1* [MIM:142200], *NOS1* [MIM 163731] and *ARG2* [MIM 107830]). Notably, although a set of variants were found in common in both good responders and poor responders across five targeted loci (*HAO2* [MIM: 605176], *BCL11A* [MIM: 606557], *WAC* [MIM: 615049], *CYP2C9* [MIM: 601130], and *CYP2E1* [MIN: 124040]), they occurred with different numbers and types of SNPs (**Figure 2A**).

**Table 1.**
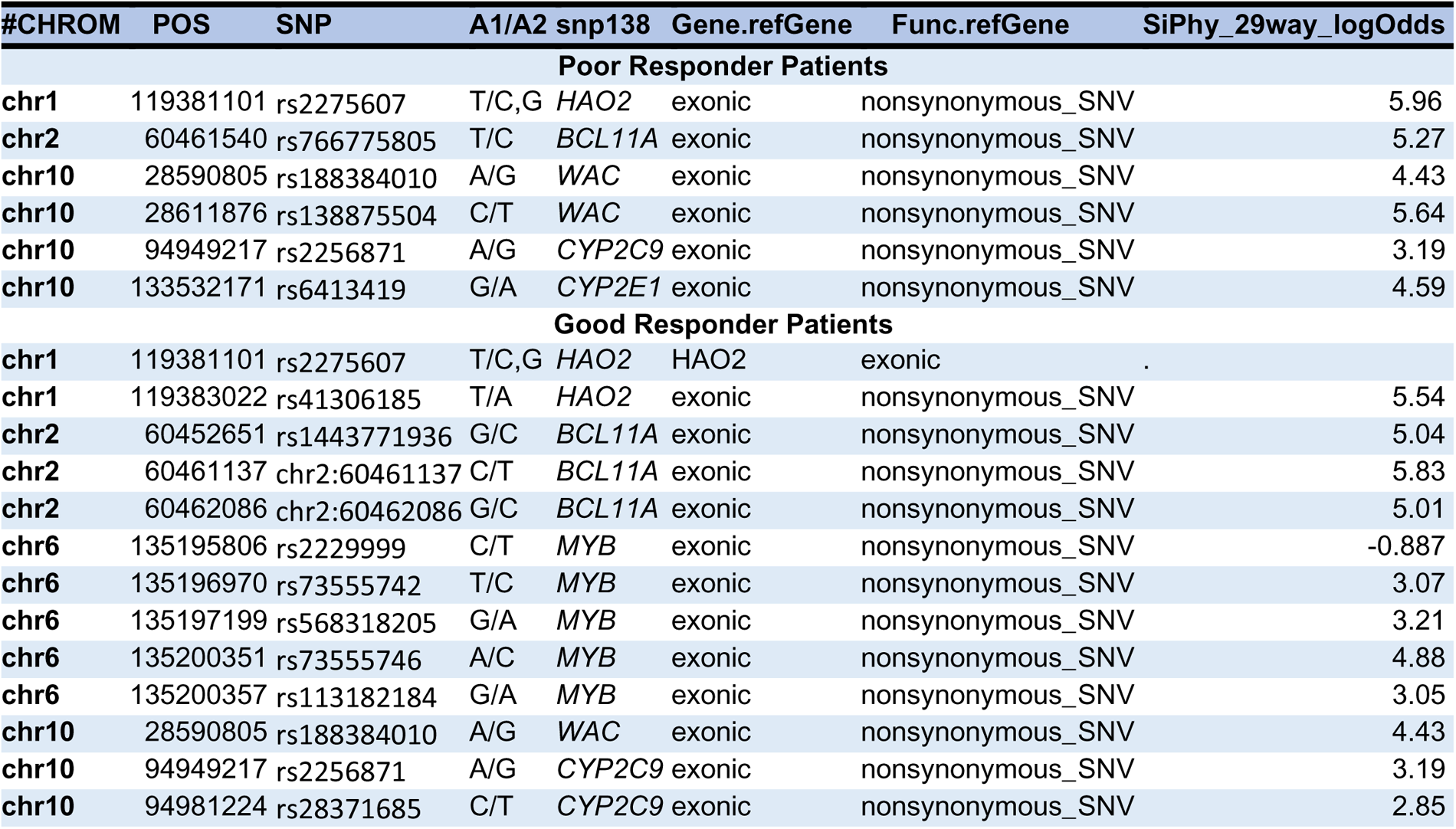
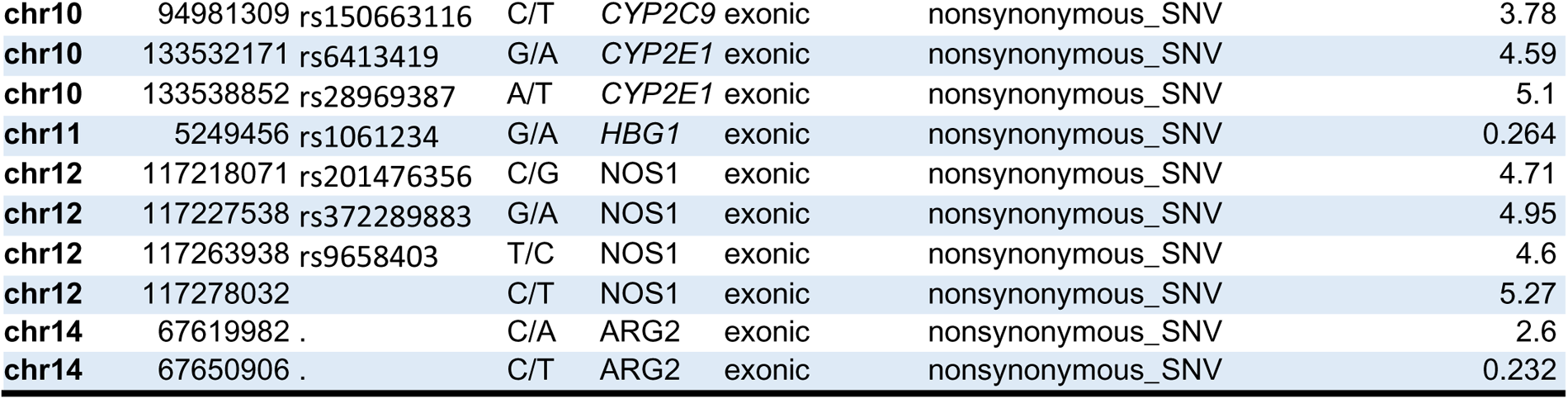
The most deleterious nonsynonymous single nucleotide variants. Exonic. nonsynonymous variants that were considered damaging according to 21 different functional scores from the annotation databases, including SIFT, LRT, MutationTaster, MutationAssessor, FATHMM, fathmm-MKL, RadialSVM, LR, PROVEAN, MetaSVM, MetaLR, CADD, GERP++, DANN, M-CAP, Eigen, GenoCanyon, Polyphen2 HVAR, Polyphen2 HDIV, PhyloP, and SiPhy.

| #CHROM | POS | SNP | A1/A2 | snp138 | Gene.refGene | Func.refGene | SiPhy_29way_logOdds |
| --- | --- | --- | --- | --- | --- | --- | --- |
| **Poor Responder Patients** |  |  |  |  |  |  |  |
| chr1 | 119381101 | rs2275607 | T/C,G | *HAO2* | exonic | nonsynonymous_SNV | 5.96 |
| chr2 | 60461540 | rs766775805 | T/C | *BCL11A* | exonic | nonsynonymous_SNV | 5.27 |
| chr10 | 28590805 | rs188384010 | A/G | *WAC* | exonic | nonsynonymous_SNV | 4.43 |
| chr10 | 28611876 | rs138875504 | C/T | *WAC* | exonic | nonsynonymous_SNV | 5.64 |
| chr10 | 94949217 | rs2256871 | A/G | *CYP2C9* | exonic | nonsynonymous_SNV | 3.19 |
| chr10 | 133532171 | rs6413419 | G/A | *CYP2E1* | exonic | nonsynonymous_SNV | 4.59 |
| **Good Responder Patients** |  |  |  |  |  |  |  |
| chr1 | 119381101 | rs2275607 | T/C,G | *HAO2* | *HAO2* | exonic | . |
| chr1 | 119383022 | rs41306185 | T/A | *HAO2* | exonic | nonsynonymous_SNV | 5.54 |
| chr2 | 60452651 | rs1443771936 | G/C | *BCL11A* | exonic | nonsynonymous_SNV | 5.04 |
| chr2 | 60461137 | chr2:60461137 | C/T | *BCL11A* | exonic | nonsynonymous_SNV | 5.83 |
| chr2 | 60462086 | chr2:60462086 | G/C | *BCL11A* | exonic | nonsynonymous_SNV | 5.01 |
| chr6 | 135195806 | rs2229999 | C/T | *MYB* | exonic | nonsynonymous_SNV | -0.887 |
| chr6 | 135196970 | rs73555742 | T/C | *MYB* | exonic | nonsynonymous_SNV | 3.07 |
| chr6 | 135197199 | rs568318205 | G/A | *MYB* | exonic | nonsynonymous_SNV | 3.21 |
| chr6 | 135200351 | rs73555746 | A/C | *MYB* | exonic | nonsynonymous_SNV | 4.88 |
| chr6 | 135200357 | rs113182184 | G/A | *MYB* | exonic | nonsynonymous_SNV | 3.05 |
| chr10 | 28590805 | rs188384010 | A/G | *WAC* | exonic | nonsynonymous_SNV | 4.43 |
| chr10 | 94949217 | rs2256871 | A/G | *CYP2C9* | exonic | nonsynonymous_SNV | 3.19 |
| chr10 | 94981224 | rs28371685 | C/T | *CYP2C9* | exonic | nonsynonymous_SNV | 2.85 |
| **chr10** | 94981309 | rs150663116 | C/T | *CYP2C9* | exonic | nonsynonymous_SNV | 3.78 |
| **chr10** | 133532171 | rs6413419 | G/A | *CYP2E1* | exonic | nonsynonymous_SNV | 4.59 |
| **chr10** | 133538852 | rs28969387 | A/T | *CYP2E1* | exonic | nonsynonymous_SNV | 5.1 |
| **chr11** | 5249456 | rs1061234 | G/A | *HBG1* | exonic | nonsynonymous_SNV | 0.264 |
| **chr12** | 117218071 | rs201476356 | C/G | *NOS1* | exonic | nonsynonymous_SNV | 4.71 |
| **chr12** | 117227538 | rs372289883 | G/A | *NOS1* | exonic | nonsynonymous_SNV | 4.95 |
| **chr12** | 117263938 | rs9658403 | T/C | *NOS1* | exonic | nonsynonymous_SNV | 4.6 |
| **chr12** | 117278032 |  | C/T | *NOS1* | exonic | nonsynonymous_SNV | 5.27 |
| **chr14** | 67619982 | . | C/A | *ARG2* | exonic | nonsynonymous_SNV | 2.6 |
| **chr14** | 67650906 | . | C/T | *ARG2* | exonic | nonsynonymous_SNV | 0.232 |

We evaluated the distribution of pathogenic SNPs situated in the targeted 13 variants in poor responders, good responders and among the general African ancestry populations (see **Materials and Methods**) (**Supplemental_Table_2B**). A considerable variation in the proportions of pathogenic SNPs across *MYB* [MIM: 189990], *HAO2* [MIM: 605176], *CYP2C9* [MIM: 601130], *ARG2* [MIM: 107830], *HBB* [MIM: 141900], *SAR1A* [MIM: 607691], *HBG2* [MIM: 142250], *KLF10* [MIM: 601878] and *CYP2E1* [MIN: 124040]) (**Table 1**), were discerned between individuals exhibiting varied responses to Hydroxyurea therapy in the context of sickle cell disease. The investigation into pathogenic SNPs across the 13 targeted variants revealed intricate patterns associated with hydroxyurea treatment response. In addition, noteworthy, *NOS1* [MIM 163731], *BCL11A* [MIM: 606557], *WAC* [MIM: 615049] and *HBG1 [MIM:142200]* demonstrated an equitable distribution of pathogenic SNPs between non-responders and responders, respectively (**Figure 2A, Supplemental_Table_2A**). In contrast to poor responders, the good responders tended to have a lower proportion of pathogenic SNPs in the targeted 13 variants, except in *CYP2C9* which showed a higher proportion. Similarly, in the general African ancestry population, a higher variation of pathogenic SNPs was observed at four (*CYP2C9, CYP2E1, HAO2* and *ARG2*), of the targeted variants (**Figure 2B, Supplemental_Table_2B**). This finding suggests that potential cumulative mutations at these genes *CYP2C9* [MIM: 601130], *CYP2E1* [MIN: 124040]*, HAO2* [MIM: 605176] and *ARG2* [MIM 107830] might be beneficial for a good response to Hydroxyurea therapy in the context of sickle cell disease.

**Figure 2.**
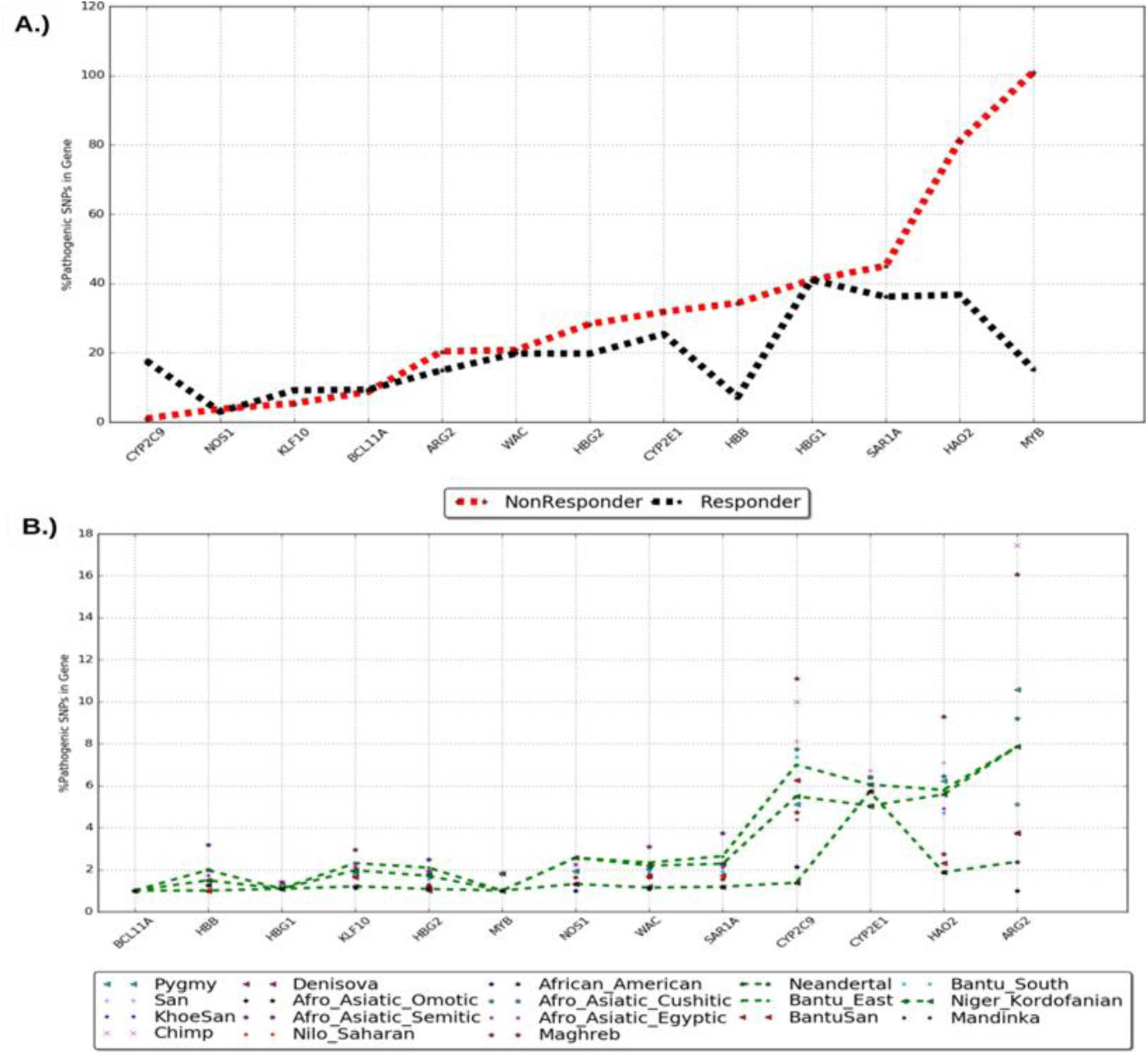
Distribution of pathogenic SNPs across the targeted genetic variants in the studied population and other ethnic groups. (A.) The upper graph illustrates the variation in the proportions of pathogenic SNPs across genetic variants among responders (black dotted line) and non-responders (black dotted line). B.) The lower graph shows proportions of pathogenic SNPs for different ethnic groups, with each group represented by a distinct colour dotted line.

### Population structure underlying the hydroxyurea specific targeted variants panel

To understand the genetic structure and diversity with respect to the identified variants (**Supplemental_Table_2A)**, Principal Component Analysis (PCA) was executed on two datasets, encompassing the general population from African ancestry descent (**Supplemental_Table_1A**), and the cohort of poor and good hydroxyurea responders from Tanzania. The top 10 principal components were retained out of 4591 (our cohort and other African populations) and 59 (only our cohort), respectively. We have plotted the distribution of these top 10 PCs (**see Supplemental_Fig_3**). Importantly, we have also indicated the eigenvalue_k/(Sum of eigenvalues) which is the proportion of variance explained by eigenvector k (https://github.com/chrchang/eigensoft/blob/master/POPGEN/README). An identifiable genetic variation pattern linked with hydroxyurea response emerged within the general population, as illustrated in **Figure 3A**. The populations represented in the PCA analysis included those in **Supplemental_Table_1A** from various other African ethnicities as well as Chimp, Neandertal and Danisova. The results in **Figure 3A** suggest that individual genetic variation with respect to the identified variants might contribute to possible variations in hydroxyurea treatment therapy. This is also supported by the assessment of genetic structure and relationships between good responders and poor responders which has revealed a discernible systematic SNPs variation among good responders, contrasted with a more homogeneous variation within the poor responder subgroup (**Figure 3B**).

### Distribution of minor allele frequencies of SNPs situated at hydroxyurea-specific targeted genes

To highlight the genetic diversity of the targeted sequenced regions in the good responders and poor responders as well as in the general African ancestry population **Supplemental_Table_1B**, we visualized binned MAF proportions, showing the distribution within predefined MAF ranges (**Materials and Methods**). The variation was observed in the distribution of MAF of rare and common variants between good responders and poor responders (**Supplemental_Fig_4A**), and a clear variation of rare variants across the rest of Africana ancestry populations and Neandertal and Maghreb (**Supplemental_Fig_4B**).

This is not surprising and might support the hypothesis of multiple rare variants associated with the SCD genetic structure among African ancestry populations, justifying the possibility of a higher prevalence of SCD in African ancestry compared to other populations. In addition, the higher variation of rare variants observed in poor responders (**Supplemental_Fig_4A**) is supported by the observed higher proportion of pathogenic SNPs (**Supplemental_Fig_4A**) in the same group compared to those found in good responders.

**Figure 3.**
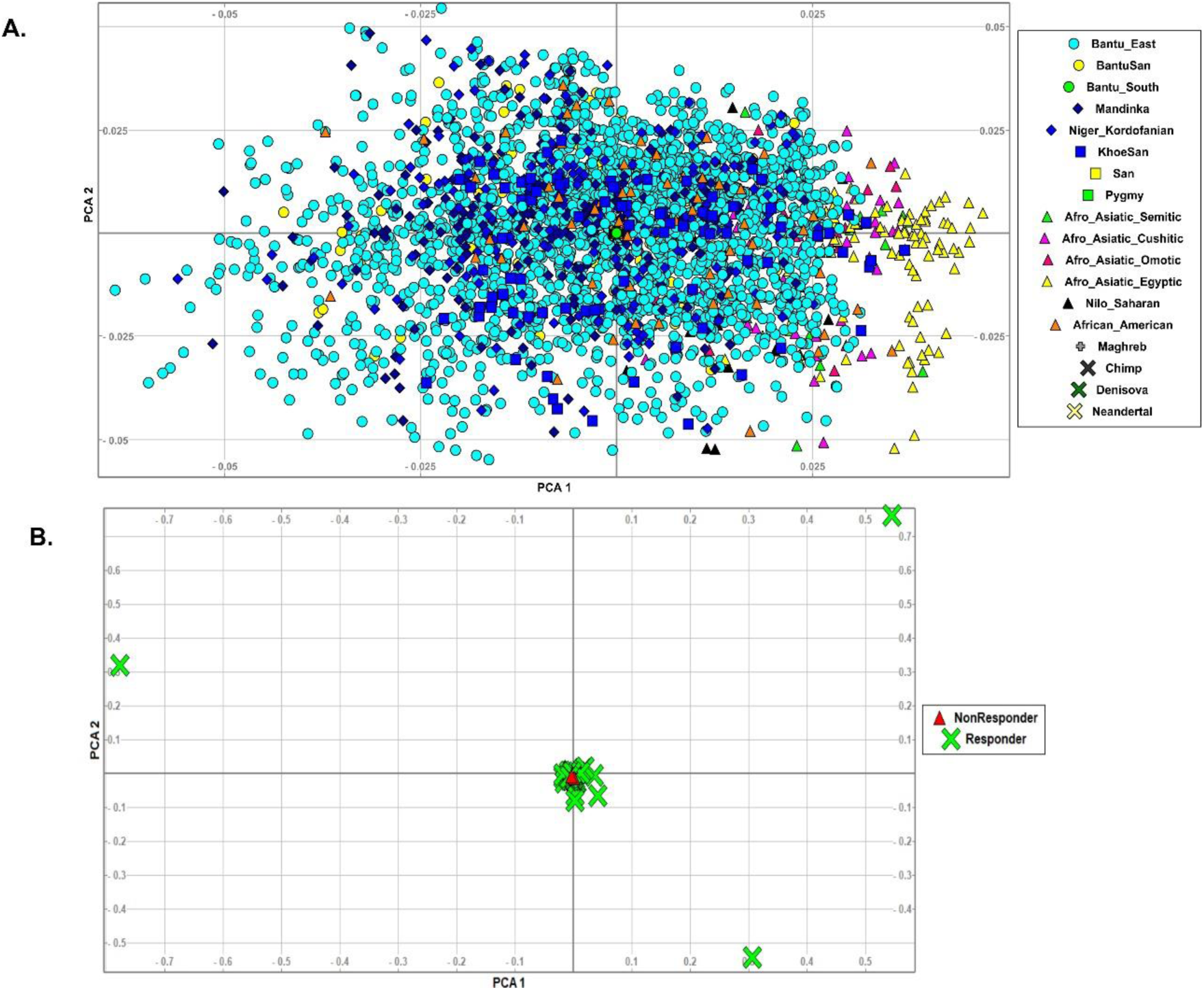
PCA plot of Hydroxyurea associated population structure across world-wide ethnicities. The diagram shows how different ethnic groups around the world are genetically related to Hydroxyurea associated genetic variants. **Plot A** compares a group of people who have specific genetic variants related to Hydroxyurea treatment with people of African ancestry from various regions, such as Bantu East, BantuSan, Bantu South, KhoeSan, San and others. The symbols on the plot represent each ethnic group, as explained by the key on the right. The plot shows that the genetic variants associated with Hydroxyurea treatment response are spread among different ethnic groups, but those found in the good responders group are more concentrated at the center. **Plot B** zooms in on the clustering of the genetic variants haboured by the good responders and poor responder groups, who are marked by green “X” and green asterisk symbols respectively. The plot shows that the responders are clustered together. This suggests that the responders share more genetic similarities with each other than with the non-responders.

The variation of SNP frequencies across the 13 targeted variants was observed in the poor responders, good responders and among the general African ancestry populations (see **Materials and Methods**) (**Supplemental_Table_1B**). A considerable variation in SNPs allele frequency (**Figure 2A**) across loci *MYB* [MIM: 189990], *HBG1* [MIM:142200], *NOS1* [MIM 163731]), *CYP2C9* [MIM: 601130], *SAR1A* [MIM: 607691] and *KLF10* [MIM: 601878] listed in **Table 1** and **Supplemental_Table_2A** were discerned between individuals exhibiting varied responses to Hydroxyurea therapy, suggesting that these variants may potentially harbour different levels of effects on treatment responses among SCD patients (**Figure 2A**, **Supplemental_Table_3A**). Noteworthy, *CYP2E1* [MIN: 124040], *(40%) and HAO2* [MIM: 605176] *(approx. 30%)* demonstrated an equitable distribution of SNPs allele frequencies between poor responders and good responders, respectively (**Figure 2A**, **Supplemental_Table_3A**). Interestingly, the good responders in contrast to the poor responders were found to have a higher proportion of SNPs across the targeted variants associated with Hydroxyurea treatment response (**Figure 2A**, **Supplemental_Table_3A**). Similar variation of SNPs frequency at these relevant genes associated with Hydroxyurea were observed in the general African ancestry populations (**Figure 2B, Supplemental_Table_3B**). This suggests individual genetic variation, or potential cumulative mutations situated on these variants may impact response to Hydroxyurea therapy in the context of sickle cell disease.

### Pathways enrichment analysis and gene-gene interactions

The 9 loci (**Table 1**) harbouring the potentially pathogenic variants were subjected to enrichment analysis using GeneMANIA and Enrichr bioinformatics tools (**Materials and Methods**) to identify molecular functions, pathways and drugs putatively affected (**Figure 4, Supplemental_Table_4**). To successfully enrich for molecular functions and pathways, the identified genes were used to “fish” more related genes that are predicted to physical and co-express interactions with the identified genes and reconstructed their network in **Figure 4**.

The products of the identified genes were predicted to perform the following predicted molecular functions (**Figure 4** and **Supplemental_Table_4A**) including tetrapyrrole binding (pvalue = 2.68 x 10^-21^), oxidoreductase activity (pvalue = 1.24 x 10^-18^), drug catabolic process and response to xenobiotic stimulus (pvalue =1.61 x 10^-07^). These gene products in **Figure 4** are mostly localized within the drug metabolism (pvalue = 4.74 x 10^-008^), metabolism of xenobiotics by cytochrome P450 (pvalue = 1.41 x 10^-007^), Arginine and proline metabolism (pvalue = 1.85 x 10^-05^), long-term depression (pvalue = 3.82 x 10^-05^), Arginine biosynthesis (pvalue = 4.40 x 10^-05^) pathways as well as associated with drugs such as Stanozolol (pvalue = 1.03 x 10^-06^) and Phenelzine (pvalue = 6.42 x 10^-06^) (**Supplemental_Table_4**).

The constructed network of genes within **Figure 4** were found to be associated with several GETex Aging Signature and tissue-specific impacting aging and genetic (**Supplemental_Table_4B**) include GTEx Blood and Thyroid (pvalue < 1.14 x 10^-06^). Importantly, our findings indicated that the constructed network of our 9 variants (**Figure 4**) is associated with TF1 haematopoietic and lymphoid tissue TenPx37 (pvalue = 1.78 x 10^-005^), supporting published transcriptomic evidence providing functional support for the role of the identified pathways (**Supplemental_Table_5**), including the arginine and proline metabolism, long-term depression, arginine biosynthesis, and metabolism of xenobiotics by cytochrome P450. This suggests that some of the identified genes may act together in these biological pathways to have a cumulative effect or influence Hydroxyurea treatment response.

**Figure 4.**
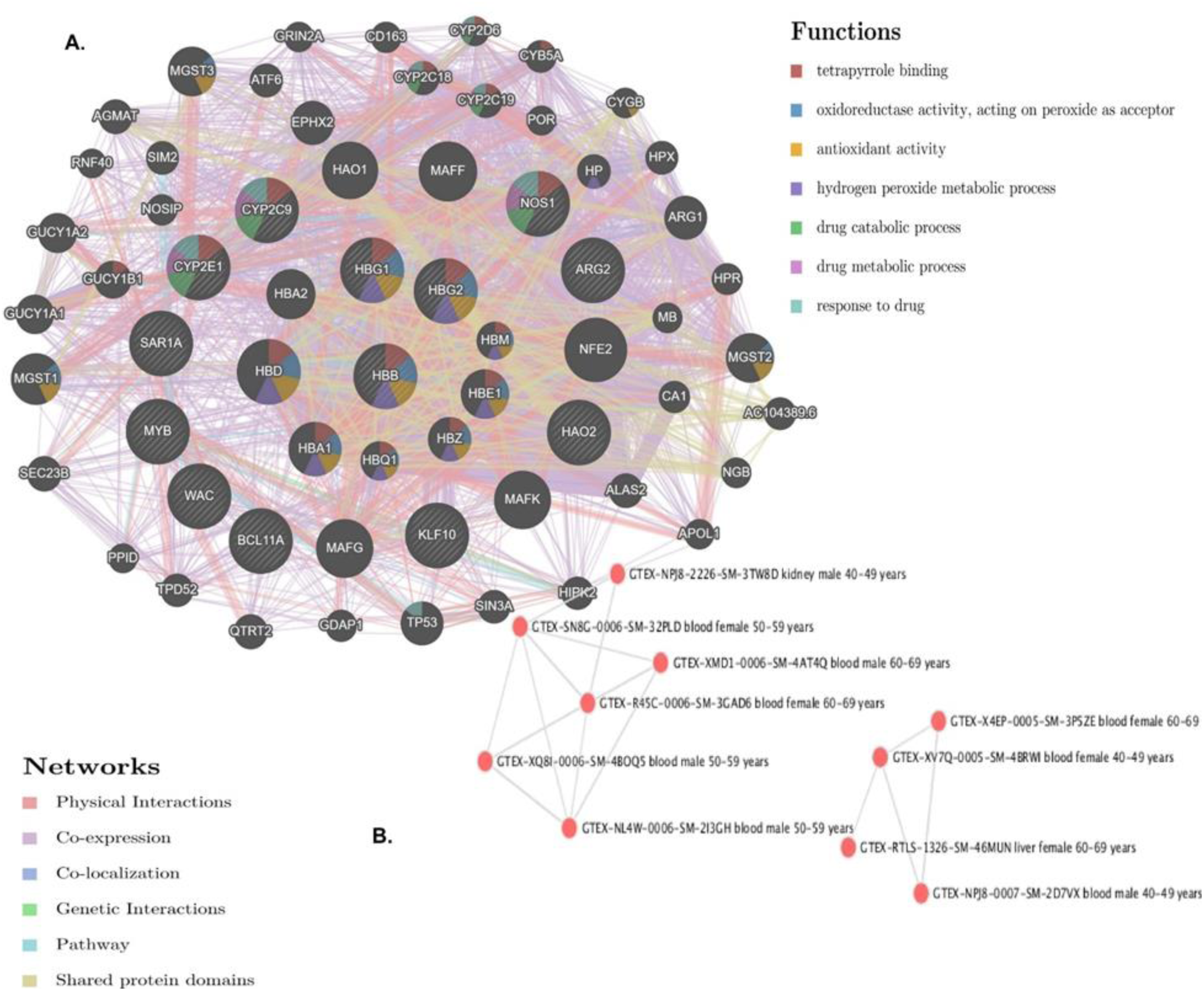
Gene-gene interaction network of genes harbouring the most deleterious variants. This figure shows how genes with deleterious variants interact with each other and with other genes. Different colours of branches represent how the genes are related; pink: physical interactions, purple: co-expression, orange: predicted, navy blue: co-localization, blue: Pathway, green: Genetic interactions, yellow: shared protein domains. Black and stripped nodes: genes provided as input into GeneMANIA. Black only nodes: genes predicted by GeneMANIA to interact with the input list. Connecting lines represent interactions. The functions of the genes are listed on the right side of the figure. The bottom part of the figure shows how samples from different age groups and genders that are linked to the genes.

## DISCUSSION

This study constitutes a groundbreaking endeavour to explore the genetic factors influencing the response to hydroxyurea (HU) in the context of Africa, where the burden of sickle cell disease (SCD) is particularly pronounced. Hydroxyurea remains the predominant disease-modifying drug for SCD globally including Africa countries such as Tanzania. Following accumulating evidence supporting the efficacy of hydroxyurea, several countries, including those in Africa, institute the administration of hydroxyurea at the age of 9 months [2,52]. The same regimen is followed in Tanzania. Notwithstanding the favourable success rates linked to the utilization of hydroxyurea, a subset of individuals with sickle cell disease (SCD) exhibits inadequate responses to the drug [53] . In line with this, our findings have also shown a proportion of patients (n=60) with poor response, particularly in relation to the levels of HbF (7.4%). Suboptimal adherence to treatment and drug dosage have been proposed as significant determinants of hydroxyurea response [53]. However, genetic factors are anticipated to contribute to this phenomenon due to their influence on drug metabolism [54]. Moreover, in the context of hydroxyurea, treatment response is notably affected by baseline levels of fetal hemoglobin (HbF) which are influenced by genetic factors across multiple loci [54]. As a result, both genetic factors that affect baseline HbF levels and those associated with drug metabolism have an impact on hydroxyurea treatment. This work built on these facts, and hence the investigation of genetic variants at 13 loci; two loci associated with hydroxyurea metabolism (*CYP2C9 & CYP2E1)* and eleven loci associated with HbF induction *(MYB, HBB, HBG1, WAC, HBG2, BCLL1A, KLF10, HA02, NOS1, ARG2* and *SAR1A)* in our SCD patient population under hydroxyurea treatment. We have also looked at hematological parameters following hydroxyurea use. Ideally, hydroxyurea response can be measured using hematological, molecular, or genetic markers **(Supplemental_Fig_1**). Of the three, genetic markers can be applied as the earliest predictors of treatment response and can be identified quite early on in life. Hence, genetic markers pose an enormous potential for development of pharmacogenomics panel for hydroxyurea treatment. Of recent, we have observed a significant increase in the development and implementation of alternative interventions/treatments for SCD. For example, two out of eleven gene-based therapies have received approval by December 2023. Similarly, more than three molecules/drugs are at different stages of trials and approvals [55–58]. The anticipated increase of this trend is poised to align with technological progress facilitating pharmaceutical development. Consequently, the utilization of early prognostic indicators, such as genetic markers associated with hydroxyurea treatment, holds significant promise in aiding informed therapeutic decisions. In addition, initial treatment decision may assist in avoiding unnecessary exposure to side effects of the drug [59].

The outcomes of our investigation have effectively fulfilled the overarching objective, namely the identification of genetic variants conducive to early prognostication of hydroxyurea treatment outcomes. Our findings distinctly indicate that while hematological parameters remain viable for monitoring hydroxyurea treatment response, their efficacy is confined to the timeframe between 3 and 6 months (refer to the results section). After monitoring the 148 individuals enrolled in the study for six months, discernible hematological profile trends were only ascertainable in those categorized as poor or good responders, as illustrated in **Figure 1**. These observations align with existing literature, underscoring the requisite at least six-month duration for the establishment of hydroxyurea response through the monitoring of hematological and other clinical profiles [2,60,61].

Looking at the genetic profiles for the patients categorised as either poor (non-responders) or good responders (responders), a remarkable difference was observed. *First*, we observed a different distribution of the prevalence of nonsynonymous single nucleotide polymorphisms across the targeted variants. For example, out of the 13 targeted loci, variants in four loci namely *MYB*, *HBG1*, *NOS1* and *ARG2* were found only in the good responders. The genetic influence of *MYB* and *HBG1* on the synthesis of fetal hemoglobin is well established in different populations including ours [62]. These findings are in line with our hypothesis that genetic loci associated with HbF levels are also associated with hydroxyurea treatment. This hypothesis is based on the established evidence that individuals with high HbF levels prior to hydroxyurea treatment respond better than those without. Similarly, as stipulated earlier, the other two loci (*NOS1* and *ARG* 2) are associated with hydroxyurea metabolism. *NOS1* is known to contribute to the conversion of hydroxyurea to its active form, which involves the release of nitric oxide [63] while Arginase 2 (*ARG2*) catalyzes the conversion of arginine, the substrate for nitric oxide synthase (NOS). Our findings are well in line with this evidence and supported by a similar trend observed for both *NOS1* and *ARG2* which are metabolically related.

Similarly, we observed different SNPs in the poor responders compared to the good responders. For example, we observed a higher number of SNPs in *CYP2C9*, *CYP2E1* and *BCL11A* in the good responders. The differential presence of SNPs at various loci may reflect the variation in hydroxyurea response across individuals. *CYP2C9* is the most abundant CYP2C subfamily enzyme which contributes to the metabolism of approximately 15% of all drugs which utilise the P450-catalyzed metabolism pathway [64]. *CYP2E1* is primarily known for its involvement in the metabolism of compounds such as ethanol and certain drugs, and its association with the generation of reactive oxygen species (ROS) and oxidative stress [65]. We observed the same for *BCL11A* which is a major HbF modifier. The association of these loci with hydroxyurea treatment responses has been reported in other studies [9,66–68], although they have not been applied for hydroxyurea response prediction in clinical settings.

*Second*, we observed differences in the prevalence of pathogenic variants between good responders and poor responders, with the latter having higher number (**Figure 2A**). Pathogenic variants are expected to cause abnormality to either gene expression or the synthesized proteins rendering them ineffective for either synthesis of HbF or hydroxyurea metabolism. Pathogenic variants may also influence hydroxyurea side effects. These findings are backed up by the variation in distribution of the variants associated with hydroxyurea treatment in the general African population (**Figure 2B**). This indicated the presence of the different variants in the population and the selection pressure that may be exerted by treatments such as hydroxyurea. These findings are augmented by what we observed from the PCA results (**Figure 3A, Supplemental_Table_2B**). The higher variation of the variants in the loci present in good responders could account for the distinct levels of hydroxyurea treatment response due to additive and interactive genetic effects. For example, we have previously reported additive effects on levels of HbF when SCD patients inherit the HbF boosting alleles at different loci [69]. It is, therefore, necessary to study in detail how genetic variants associated with hydroxyurea response interacts with one another and with other genes.

*Third*, a different distribution was observed when we looked at the minor allele frequencies (MAFs) between responders and non-responders (**Supplemental_Fig_4A**). The high frequency of rare variants in the non-responders aligns well with the high presence of pathogenic variants. Rare genetic variants have been associated with low response to drug treatment, contributing to the heritable variability in drug response. Studies have revealed that rare pharmacogenetic variants are strongly enriched in mutations predicted to cause functional alterations, and they significantly contribute to the inter-individual variability in drug response. For example, rare genetic variants have been found to influence the clinical response to anti-TNF therapy, with low-frequency loss-of-function variants being associated with the therapy’s response [70] .

*Fourth*, the SNPs allele frequencies varied across our targeted regions with a higher frequency in the responders than non-responders (except for *CYP2C9*) (**Supplemental_Fig_5A**). SNP allele frequencies provide insights into the genetic variability in drug response among different individuals and populations and hence variability in frequencies across the targeted regions. This information is vital for developing personalized medicine and optimizing drug treatments based on individual genetic profiles [66,71] .

Our study has investigated selected regions associated with either HbF synthesis or hydroxyurea metabolism. This was confirmed by the pathway’s enrichment analysis and gene-gene interactions analysis (**Figure 4**). Our findings have highlighted various predicted molecular functions including tetrapyrrole binding, oxidoreductase activity, drug catabolic process and response to xenobiotic stimulus and mostly localized within the drug metabolism such as metabolism of xenobiotics by cytochrome P450, Arginine and proline metabolism, Arginine biosynthesis pathways. These functions and pathways have been previously associated with drugs metabolism including hydroxyurea [72–74]. These findings are in line with the genetic profiles we have described for responders and non responders. For example, the metabolic pathway involving *NOS1,* and hydroxyurea is intricate and involves the enzymatic conversion of hydroxyurea to nitric oxide. In addition, looking at the pathways and interactions between the targeted genes/loci and other gives us an opportunity to identify additional potential drug targets for SCD.

## Materials and Methods

### Patient selection and enrolment into the study

The study included a total of 150 SCD patients followed up for six months at Muhimbili National Hospital (MNH), Amana and Temeke hospitals. The study selectively enrolled patients aged five years and above who had not previously received hydroxyurea treatment and met specific haematological criteria for hydroxyurea initiation: HbA <15%, absolute neutrophil count >2,000/uL, platelets >100,000/uL, hemoglobin level >5.0g/dL, and absolute reticulocyte count >100,000/uL. Additionally, comprehensive anamnesis was conducted at the time of enrolment, encompassing an inquiry into prior disease manifestations and concurrent symptoms. The research excluded pregnant and lactating women, individuals planning conception during the study period, and those undergoing chronic transfusion therapy.

### Hydroxyurea treatment and monitoring

Each patient was subjected to clinical examination by a specialised haematologist and collection of a whole blood sample for haematological analysis, HbF quantification and DNA extraction. Complete blood count (RBC, reticulocytes, WBC, and platelets), haemoglobin, MCV, and MCH were determined using the XT-2000i Haematology Analyzer (Sysmex Corporation, Kobe, Japan).

The haemoglobin fractions, including HbF, were quantified by HPLC (Bio-Rad Variant II, Hercules, CA, USA). The eligible patients started the treatment with HU (daily dose 20 mg/kg), and every month had a follow-up consultation, where all the manifestations and symptoms were registered.

The assessment of the response to hydroxyurea (HU) treatment hinged on alterations in fetal hemoglobin (HbF) levels, leading to the classification of participants into responders and non-/poor responders. Responders were identified as patients exhibiting a two-fold or greater increase in HbF levels compared to baseline which could be easily captured within three to six months post-treatment. The evaluation of compliance to hydroxyurea treatment encompassed analyses of haematological parameters, specifically an augmentation in total hemoglobin, mean corpuscular volume (MCV), and mean corpuscular hemoglobin (MCH). Compliance was further gauged through pill counting and self-reported adherence. This data was instrumental in the identification and exclusion of non-compliant participants from the non-/poor responder’s cohort. Out of the 150 participants, 48 participants were categorized as extreme good responders (greater than two-fold increase in HbF levels) while 11 were extreme poor responders (same levels of HbF compared to the baseline levels). The rest of the participants were categorized as responders (two-fold increase in HbF levels).

### DNA extraction, library preparation and sequencing

Genomic DNA was extracted and purified from peripheral blood samples using the QIAamp DNA Blood Mini Kit (Qiagen GmbH, Hilden, Germany) according to the manufacturer’s recommendations. All samples were quantified by a Qubit™ dsDNA HS fluorometric assay (ThermoFisher Scientific Inc., Waltham, MA, USA).

A custom Ampliseq panel was used for interrogating genetic variants the 13 loci (**Supplemental_Table_1B**). The libraries were prepared with Ampliseq Library Plus for Illumina kit (Illumina, San Diego, CA, USA and using Illumina’s Ampliseq CD indexes set A(Illumina), and their quality was assessed with High Sensitivity DNA reagents on an Agilent 2100 Bioanalyzer System. Paired-end sequencing was performed on a MiSeq sequencer (Illumina, San Diego, CA, USA) using the MiSeq Reagent kit v3 (600 cycles) as in our previous paper [12].

The current study scrutinized the intricate genetic milieu of genes linked to hydroxyurea response within a cohort comprising 11 non-responders and 49 responders among sickle cell disease (SCD) patients with hemoglobin F (HbF) in Tanzania. Additionally, the analysis was extended to encompass ethnolinguistic groups of general African descent.

### Data management and statistical analysis

#### Haematological profiling

A linear mixed model analysis was conducted on a sample of 60 patients to evaluate the impact of hydroxyurea treatment on four key haematological indices: Hemoglobin F (HBF), Mean Corpuscular Volume (MCV), White Blood Cell count (WBC), and Reticulocyte count (RET).

In this model, both the number of visits and the response status (responder or non-responder to treatment) were included as fixed effects. The response status was treated as a moderator to examine if the effect of the number of visits on the haematological indices differed between responders and non-responders. To assess this moderation effect, product terms were included in the model. The model specification was represented as follows for each dependent variable (haematological index): Dependent Variable∼Visits+Responding Status+Visits:Responding Status+(1∣Subject). This specification indicates that the dependent variable (each haematological index) is modelled as a function of the number of visits, the responding status, and the interaction between visits and responding status, with a random intercept for each subject to account for within-subject correlation.

The variable “visits” in the model is a count variable representing the number of visits each patient had during the study period. The random-effect term is assumed to follow a normal distribution, which is a common assumption in linear mixed models. To control for potential confounding effects, age and gender were included as covariates in the model. These variables were chosen because they are known to influence the haematological indices and could potentially confound the relationship between hydroxyurea treatment and these indices. Before conducting the analysis, the assumptions underlying the linear mixed-effects model were assessed. This included checking for normality and homogeneity of variance of residuals, sphericity, and linearity for quantitative predictors.

#### Variant Calling and Downstream Data Description

After performing targeted sequencing, the quality of the sequencing reads was assessed before further analysis. FastQC [13] and SolexaQC++ [14] were used for quality control. The sequencing reads were then aligned to the human reference genome build grch38.p2 obtained from the UCSC database [15] using BWA-MEM [16,17] with local realignment conducted using the Genome Analysis Toolkit (GATK) software [18]. Additional FastQC analysis was performed on the final BAM files before variant calling, and MultiQC [19] employed to consolidate the FastQC results into a single report. All sequencing sequences passed the quality control criteria.

Joint variant calling was performed on sixty subjects from Tanzania (including 11 non-responders and 49 responders) with HbF patients treated with hydroxyurea. The variant calling was done using GATK [18,20] and BCFTOOLS [21]. To enhance the accuracy of variant calls, VariantMetaCaller [22] was employed, a support vector machines approach, which combined and optimized the variant calls based on the statistical properties and discovery consensus from both BCFTOOLS and GATK callers. The resulting variant calling files were filtered using the GATK tool “Variant Filtration”. We reported the overall quality of data in **Supplemental_Fig_2.**

#### Description of other sequence data used

To elucidate the genetic underpinning hydroxyurea response in general African, descend populations, we leveraged our previously merged data from 1000 Genomes Project Consortium, 2015 [23] and the African Genome Variation Project (AGVP) [24]. From this dataset, we only use the samples from African descend population comprises 9,578,615 Single Nucleotide Polymorphisms (SNPs) meticulously curated from the dbSNP database [25]. We utilized ethnolinguistic markers to categorize individuals, resulting in the precise classification into eighteen (18) distinct African descend groups. **Supplemental_Table_1A** offers a comprehensive look at the demographic details of these groups, encapsulating a broad representation of global African descend populations (African American, Afro_Asiatic_Cushitic, Afro_Asiatic_Egyptic, Afro_Asiatic_Omotic, Afro_Asiatic_Semitic, Bantu_East, BantuSan, Bantu_South, KhoeSan, KhoeSan, Maghreb, Mandinka, Niger_Kordofanian, Nilo_Saharan, Pygmy and San. We centred our efforts on extracting Single Nucleotide Polymorphisms (SNP) data aligning to our selected set of hydroxyurea-associated genes. The dbSNP database [25], with its extensive repositories, paired with our gene list, formed the backbone of this extraction. We strategically queried the dbSNP database using identifiers from our gene compilation, pinpointing SNPs directly associated with these hydroxyurea-specific genes. This process produced an expansive set of 63,516 SNPs, intensifying the granularity of our subsequent analysis focused on genetic variations of hydroxyurea response among SCD patients. We leveraged this set of SNPs to perform hydroxyurea-specific population structure, distribution of allele frequency and of pathogenic at these SNPs associated to hydroxyurea-specific genes (see below sections).

#### Variant annotation and mutation prioritization

The resulting VCF files were annotated using ANNOVAR [26] and independently performed variant-based annotation in each final VCF data set to determine whether SNPs cause protein-coding changes and produce a list of the amino acids that were affected. Population frequency, pathogenicity for each variant from 1000 Genomes data [23], Exome Aggregation Consortium (ExAC) [27], targeted exon datasets and COSMIC [28] were obtained with ANNOVAR database settings [[26]. ANNOVAR’s library and RefGene [29] were used to extract variant function and different functional predictions. ANNOVAR has at least up to 21 different mutation score tools including SIFT[30], LRT[31], MutationTaster [32], MutationAssessor[33], FATHMM and FATHMM-MK[34,35], RadialSVM[36], LR[36], PROVEAN[36], MetaSVM[36], MetaLR[36], CADD[36,37], GERP++[38], DANN[39], M-CAP[40], Eigen[41], GenoCanyon[41], Polyphen2-HVAR [42]and HDIV[42], PhyloP[43], and SiPhy[43]. In addition, conservative and segmental duplication sites were included, dbSNP code and clinical relevance reported in dbSNP[25]. From the resulting functional annotated data set, we independently filtered for predicted functional status (of which each predicted functional status is of “deleterious”(D), “probably damaging” (D), “disease-causing-automatic” (A) or “disease-causing” (D). The candidate mutation was selected based on step 1) casting vote approach implemented in our custom python script, to retaining only a variant if it had at least 17 predicted functional status “D” or “A” out of 21 was used and step 2) further filtering for rarity, exonic variants, and nonsynonymous mutations and with a high-quality call from the retained variants from step 1 above.

#### Hydroxyurea-associated pathogenic SNPs

We then moved to investigate the distribution of pathogenic Single Nucleotide Polymorphisms (SNPs) that are specifically associated with hydroxyurea treatment response variation among SCD and across general African descent ethnic populations. This phase of our analysis involved meticulously grouping genetic data, enriched with details about pathogenic SNPs. This stratification allows us to calculate the proportions of pathogenic SNPs (predicted as described in section ***Variant annotation and mutation prioritization*)** within each group. Using Python programming along with specialized libraries such as “matplotlib” and “NumPy”, visual representations were crafted that spotlighted the ethnic-specific proportions of pathogenic SNPs for individual genes. The genetic data, combined with information about pathogenic SNPs and their associations with different ethnic groups, is systematically organized according to group.

#### Population structure associated with hydroxyurea treatment response

To evaluate the extent of substructure within our SCD cohort, the curated 60 phased haplotypes dataset, resulting from Eagle [44] was leveraged to perform genetic structure analysis based on Principal Component Analysis (PCA). Furthermore, a PCA analysis was performed to investigate the genetic structure based on 63,516 SNPs African descent population groups. A quality control check was conducted on these VCF files using plink [45]. Principal Component Analysis (PCA) via EIGENSOFT 3.0 package’s smartpca tool [46,47] was employed using the two datasets as a powerful tool to unravel underlying substructures within hydroxyurea-related polymorphisms (**Supplemental_Fig_3**). This analysis enabled the distillation of complex genetic variations into a reduced-dimensional space. The PCA plots generated were subsequently visualized using the Genesis software [48], allowing us to gain insightful perspectives on the genetic diversity and variations associated with hydroxyurea pharmacogenomics within these populations.

#### Distribution and frequencies of SNPs across targeted variants associated with hydroxyurea response

The distribution of the minor allele frequency for the identified SNPs situated in the 13 targeted genetic variants were investigated in order to understand which SNPs are common and those which were rare. The proportion of minor alleles frequencies were categorized into 6 bins (0-0.05, >0.05-0.1, >0.1-0.2, >0.2-0.3, >0.3-0.4, >0.4-0.5) with respect to each ethnic group as well as in our hydroxyurea responder versus non-responder patients. The minor allele frequency (MAF) per SNP for each category was computed using Plink software. Utilising the dbSNP database, we went on to investigate the frequency of SNPs across the targeted variants. The computation included an assumption that SNPs in upstream and downstream within a gene region are close and possibly in Linkage Disequilibrium (LD) [49].

In tandem, in order to understand the genetic diversity underlying the hydroxyurea associated SNPs across the different ethnic backgrounds, we performed a detailed analysis of the distribution of Minor Allele Frequencies (MAF) of the targeted hydroxyurea variants. We utilised a tailored Python script to process the raw MAF data, keeping in view gene annotations and a list of ethnic groups. This approach efficiently enabled us to extract MAF frequencies of the targeted hydroxyurea variants, forming an essential bedrock for our genetic exploration. Additionally, binned MAF proportions were visualised to emphasize the scope of genetic variation, delineating the distribution within predefined MAF ranges. This step involved the calculation of the proportions of genetic variations nestled within distinct MAF bin boundaries, create visual representations, crystallize trends and variances in genetic diversity across populations within unique MAF bins.

#### Pathway enrichment analysis of Hydroxyurea-associated variants

To unravel the functional implications of hydroxyurea-associated variants, enrichment analysis was conducted employing innovative tools, including GeneMania [50] and Enrichr [51]. First, we reconstructed a functional, physical and co-expression interacting network of hydroxyurea-associated variants via GeneMania [50]. This multifaceted approach enabled us to comprehensively elucidate these variants’ molecular context and potential pathways. First, we used GeneMania to explore functional networks involving the hydroxyurea-associated variants. Secondly, we examined how the constructed networks created from these variants were associated with human phenotypes, pathways, biological processes, and molecular functions using Enrichr [51]. Gene ontology terms and annotations from the Gene Ontology databases were extracted for cellular, drug components, human phenotype, and molecular functions.

## Data Access

Genomic sequencing data generated in this study have been submitted to the NCBI BioProject database (https://www.ncbi.nlm.nih.gov/bioproject/) under accession number PRJNA1114738. The original contributions presented in the study are included in the article/supplementary material, further inquiries can be directed to the corresponding authors.

## Conclusion

Our study represents a pioneering effort in investigating the genetic factors influencing the response to hydroxyurea (HU) treatment in sickle cell disease (SCD) within the context of Africa. Despite hydroxyurea’s established efficacy as a disease-modifying drug globally, including in African countries like Tanzania, a subset of individuals with SCD exhibit inadequate responses to the treatment. Our study focused on elucidating the role of genetic factors in hydroxyurea response, considering their influence on drug metabolism and baseline levels of fetal hemoglobin (HbF). Through the analysis of genetic variants at 12 loci, we identified distinct genetic profiles associated with responders and non-responders, with specific loci influencing HbF levels and hydroxyurea metabolism showing differential distribution. The identification of pathogenic variants and the observation of rare genetic variants in non-responders underscore the genetic complexity influencing hydroxyurea response. Our emphasis on the early identification of genetic markers associated with hydroxyurea treatment holds significant promise for the development of a pharmacogenomics panel, enabling informed therapeutic decisions and potentially minimizing unnecessary exposure to drug side effects. Additionally, our study contributes to the growing body of knowledge in the context of alternative interventions for SCD, aligning with the increasing development and implementation of gene-based therapies. By providing a comprehensive analysis of gene-specific SNPs, allele frequencies, and enriched molecular functions and pathways associated with hydroxyurea metabolism and HbF synthesis, our study adds depth to our understanding of the intricate genetic landscape influencing hydroxyurea treatment. The findings not only facilitate early prognostication of hydroxyurea treatment outcomes but also lay the foundation for future research aimed at unravelling the complex genetic interactions influencing drug response in SCD. This knowledge is pivotal for advancing personalized medicine approaches and optimizing drug treatments based on individual genetic profiles, thus contributing to ongoing efforts to improve outcomes for individuals with SCD.

## Limitations of our findings

The outcomes of this investigation bear substantial relevance in shaping potential alternative interventions, including the consideration of combination therapies. Furthermore, the findings may inform the development and implementation of specialized care approaches, such as alternative therapeutic molecules and gene-based therapies, to address the specific challenges posed by SCD within the African population. The study is limited in the small sample size, investigating further the identified genetic variants in order to elucidate how they influence hydroxyurea treatment.

## Ethics Statement

Ethical approval and consent to participate and publication of findings.

This study received ethical approval from the Institutional review board at Muhimbili University of Health and Allied Science, number MUHAS-REC-2018-078. Consent forms were signed by the mothers who agreed their children to participate and publishing of findings after education was given by the research assistant.

## Author Contributions

SN contributed to conceptualization, funding acquisition, supervision, project administration. CN, ERC, EB, ES, FK and HC data curation, formal analysis, investigation, methodology, and software. ERC, CN, SN and CD contributed in, writing - review and editing, conceptualization, methodology, supervision, validation, resources, visualization. DS, DN, JJ, FU, JM, SK, MZ, RS, CC, EC, EN, and JM critically reviewed the data and edited the manuscript.

## Acknowledgments

The Sickle cell Program and the Haematology Clinical and Research Laboratory of the Department of Haematology and Blood Transfusion, Muhimbili University of Health and Allied Sciences is highly appreciated for the cooperation shown during the whole time of study.

